# An end-to-end framework for single-cell-resolution, whole-transcriptomic spatial profiling in post-mortem human brain

**DOI:** 10.64898/2026.07.28.740610

**Authors:** Ayslan Castro Brant, Ekaterina Aladyeva, Hoang-Tuong Nguyen-Hao, Katharine Alltop, Nicholas Sweeney, Tae Yeon Kim, Iara D. De Souza, Subhodip Adhicary, Ricardo D ’o Albanus, Krishna L. Bharani, Hongjun Fu, Gordon Meares, Greg T. Sutherland, Oscar Harari

## Abstract

Single-cell resolution spatial transcriptomics enables transcriptome-wide molecular profiling within intact tissue architecture, providing unprecedented opportunities to investigate cellular organization and disease-associated molecular states in the human brain. However, applying these technologies to post-mortem human brain tissue remains challenging due to RNA degradation, heterogeneity in tissue preservation, and a lack of standardized analytical workflows. These challenges are particularly pronounced for whole-transcriptome platforms, where successful implementation requires optimization of both experimental and computational procedures. Here, we present an end-to-end framework for single-cell-resolution, whole-transcriptome spatial transcriptomics of fresh-frozen (FF) and formalin-fixed paraffin-embedded (FFPE) post-mortem human brain tissue. The framework combines an optimized experimental workflow with a preservation-agnostic bioinformatics pipeline for data processing, integration, and annotation. Experimentally, we show that a condensed two-day *Visium HD* workflow provides improved library quality, lowered qPCR cycle thresholds, and more consistent fragment size distributions. Sequencing saturation analyses further identified cost-effective sequencing depths that maximize transcript recovery while minimizing redundant sequencing. Computationally, we established a scalable workflow incorporating DAPI-based nuclear segmentation, transcript assignment, quality control, reference-guided integration, clustering, and cell-type annotation. We implemented a reference-based highly variable gene selection strategy to enable robust cross-sample harmonization independent of tissue preservation method. Application of this framework to seven Alzheimer’s disease frontal cortex specimens (five FF and two FFPE) generated a unified single-cell spatial transcriptomic atlas comprising more than 530,000 spatially resolved cells. The integrated dataset resolved major neuronal, glial, and vascular cell populations, recapitulated expected cortical architecture, and enabled direct comparison of FF- and FFPE-derived spatial transcriptomic profiles. Together, this work provides a practical experimental and computational framework for single-cell resolution, whole-transcriptome spatial transcriptomics in post-mortem human brain tissue and delivers a publicly available resource that expands the utility of archived and frozen specimens for studies of neurodegeneration and other neurological disorders.

**Importance:** Spatial transcriptomics of human post-mortem brain tissue is limited by RNA degradation, preservation variability, and lack of standardized workflows. Here, we present an end-to-end framework combining an optimized two-day *Visium HD* protocol with a preservation-agnostic bioinformatics pipeline. This approach improves library quality, defines efficient sequencing strategies, and enables robust spatial profiling across both FF and FFPE samples. Applied to Alzheimer’s disease brain tissue, it generates high-resolution single-cell spatial data and expands the utility of archived and frozen specimens for studying neurodegenerative diseases.

## Introduction

Spatial transcriptomics has emerged as a powerful approach for characterizing gene expression within intact tissue architecture, enabling the investigation of cellular heterogeneity, molecular interactions, and disease-specific spatial patterns at unprecedented resolution ^1^. Neurodegenerative diseases such as Alzheimer’s disease (AD) are particularly well suited for spatially resolved molecular profiling, as key neuropathological hallmarks, including amyloid-β (Aβ) plaques and neurofibrillary tangles (NFTs), occur in region-, layer-, and cell-specific patterns across the human brain ^2,3^. Understanding the transcriptional landscape at single-cell scale near pathological loci requires methods that capture RNA with high spatial fidelity in human brain tissues, including post-mortem samples.

Conventional neuropathological workflows rely primarily on FFPE tissue, which remains the gold standard for long-term preservation, histological evaluation, and diagnostic assessment ^4,5^. Although there are published guidelines for a standardized approach to collect and characterize post-mortem brain samples for neurodegenerative diseases, such as those from the National Institute on Aging and Alzheimer’s Association (NIA-AA), there still exists significant variability across NIA-AA funded brain banks in region sampling, FFPE tissue banking, and characterization ^6^. In cases where there are no clear guidelines or standardized approach (e.g, cases for normal aging, neurodevelopmental disorders, and neuropsychiatric disorders), there exists an even greater variability of available FFPE brain tissue from certain regions. This variability in collection protocols across brain banks limits harmonization between brain banks and makes it challenging to complete robust large cohort post-mortem studies. In addition, although FFPE preservation provides excellent structural integrity, nucleic acid quality is compromised by crosslinking and fragmentation, posing challenges for transcriptomic profiling ^7,8^. These limitations highlight the need for complementary preservation strategies, such as FF tissue. Although FF tissue does have inferior structural integrity compared to FFPE, it requires less processing to bank and provides better preserved RNA integrity ^7^.

Despite its potential, spatial transcriptomics using FF human post-mortem brain tissue remains technically challenging. RNA degradation due to prolonged post-mortem intervals, variability in tissue handling, and the intrinsic fragility of frozen brain tissue can compromise downstream library preparation ^7,9,10^. These challenges are further exacerbated by the complexity of commercial workflows, such as the *10X Genomics Visium HD* platform, which enables transcriptome-wide spatial profiling at the single-cell scale. The *10X Genomics Visium HD* standard protocol includes multiple optional stopping points that may introduce additional freeze–thaw cycles, thereby increasing the risk of RNA deterioration. Although the *Visium HD* system provides a standardized framework, there is limited guidance on its application to FF post-mortem human brain tissue, and no established workflow specifically optimized to mitigate the impact of processing interruptions and RNA instability in this tissue type.

To address these challenges, we evaluated and optimized key procedural steps of the *Visium HD* protocol for both FF and FFPE human post-mortem brain tissue and compared the efficiency of these preservation strategies for spatial transcriptomics. Here, we present an optimized two-day *Visium HD* workflow that minimizes processing interruptions and incorporates extended probe-hybridization. Our results demonstrate that reducing workflow pauses and increasing hybridization time significantly improve library performance, resulting in consistent fragment size distributions and reduced PCR cycle requirements. By directly addressing limitations in current workflows, enabling robust spatial transcriptomic profiling from challenging post-mortem brain samples, this study provides a practical and scalable solution to expand the application of high-resolution spatial transcriptomics in human neurodegenerative disease research.

## Methods

### Human brain tissue

Human post-mortem brains for the fresh-frozen protocol were obtained at autopsy at various sites and collected by the National Disease Research Interchange (NDRI) or the Neuroscience Research Institute Brain Bank and Biorepository (NRI-BBB) where specimens were de-identified and sent to the researcher. Samples for this study were selected based on having a clinical diagnosis of AD and a high degree of post-mortem AD Neuropathologic Change (ADNC) as indicated by ABC scores where available ^11^. To minimize potential confounding effects, we matched for age at death, sex, postmortem interval (PMI), and APOE genotype as close as possible. Following initial sample selection, RNA quality was assessed using RNA Integrity Number equivalent (RINe) scores, and those that did not meet the minimum RINe threshold required for downstream analyses were excluded and replaced with alternative samples that met quality-control criteria while maintaining comparable demographic and clinical characteristics whenever possible. Approval of this research was obtained from The Ohio State University Biomedical Institutional Review Board (2024E0190). Brains destinated for the FFPE workflow were obtained by the New South Wales Brain Tissue Research Centre (BTRC), in accordance with approval from the Scientific Advisory Committee and the University of Sydney Human Research Ethics Committee (2024/HE001724).

### Fresh-frozen human brains processing

To prepare the samples for the spatial transcriptomics workflow, a tissue fragment of approximately 0.5–0.8 cm³ was rapidly dissected and embedded in optimal cutting temperature (OCT) compound from frozen dorsolateral prefrontal cortices (DLPFC) that were stored in a −80 °C freezer. Extra care was taken to minimize tissue thawing. The OCT-embedded tissue block was then snap-frozen in an isopentane bath chilled with dry ice and subsequently stored at −80 °C until use in our two-day-optimized protocol.

Our spatial transcriptomics workflow for FF human post-mortem brain tissue was adapted from the *10X Genomics Visium HD* user guide and protocol (CG000685 and CG000763), and the previous *10X Visium SD* workflow ^12^. Prior to initiating tissue processing, all materials, instruments, and the laminar flow hood were thoroughly cleaned, sterilized, and treated with RNaseZap™ RNase Decontamination Solution (Invitrogen) to minimize the risk of RNA degradation. Because residual RNaseZap™ can interfere with downstream RNA preservation, all treated surfaces and tools were subsequently wiped with RNase-free water to remove excess reagent and then allowed to air-dry for several minutes before use.

On Day 1, RNA integrity was assessed prior to initiating the *Visium HD* workflow. First, OCT-embedded brain tissue blocks were sectioned on a cryostat, and approximately four consecutive sections (25 µm each) were collected into a 1.5 mL tube for RNA extraction using the RNeasy Kit (QIAGEN). Then, RNA integrity was quantified using the TapeStation system (Agilent), and samples with an RINe score greater than 4 were deemed suitable for downstream processing. An additional consecutive 10 µm section from samples meeting this RNA quality threshold was subsequently used for immunofluorescence (IF) staining. Optionally, an additional consecutive 10 µm section can be collected for hematoxylin and eosin (H&E) staining to assess overall tissue integrity and cellular morphology prior to downstream processing.

Tissue sections were co-stained for amyloid-β (82E1, IBL-America, catalog no. 10323), phosphorylated tau (pSer202/pThr205 – AH36, StressMarq Biosciences, catalog no. SMC-601), and DAPI (ThermoFisher) for nuclear visualization. Tissue fixation, permeabilization, and IF procedures were performed according to the *10X Genomics Visium HD* protocol. To minimize RNA degradation during staining, RNase inhibitor (Applied Biosystems) was added to the blocking buffer, primary and secondary antibody solutions, and mounting medium, as recommended in the *Visium HD* IF staining guidelines. Additionally, to reduce autofluorescence during imaging, we treated sections with 1× TrueBlack® Lipofuscin Autofluorescence Quencher (Biotium), an optional step included in the *Visium HD* protocol. Whole-tissue images were acquired on a Nikon Eclipse Ti2-E fluorescence microscope using motorized z-stacks and tile stitching at 10x magnification. Immediately after IF image acquisition, the tissue slides were decoverslipped, and the probes were hybridized for 20 hours.

On Day 2, after completion of the probe-hybridization incubation, all subsequent steps were performed consecutively without using any of the optional stop points recommended in the original *Visium HD* protocol. The second day of the protocol began with the post-hybridization washes, followed by probe ligation and the corresponding post-ligation wash steps. During preparation of the *Visium HD* slides for subsequent stages, tissue-containing slides were kept in a thermocycler at 4 °C to preserve RNA integrity. Probe release, extension, and capture using the CytAssist instrument were performed according to the manufacturer’s user guide, again without incorporating any stop points. After elution, probes were immediately subjected to pre-amplification and SPRIselect cleanup, followed directly by library construction. Libraries were generated following the *10X Genomics* recommendations, and final cleanup was performed using SPRIselect beads. Post-library-construction quality control (QC) was assessed using the TapeStation D1000 ScreenTape system (Agilent). Libraries were subsequently sequenced in one line of a 10B flow cell using paired-end 150 bp reads (PE150) on a NovaSeq X Plus (Illumina). In total, the protocol required approximately 8 hours of hands-on time on Day 1 and approximately 12 hours on Day 2, in addition to a 20-hour incubation for probe hybridization.

### FFPE human brains processing

Human post-mortem brains were collected at autopsy and hemisected for both fresh-frozen and FFPE processing. For the latter, the hemisphere was initially perfused with 15% neutral buffered formalin (NBF) for 1 hour, then cut into 0.5-cm-thick blocks and further immersed in 10% NBF for 24 hours. Blocks were subsequently processed using the Excelsior AS (Epredia) automated tissue processor, where dehydration was carried out with graded ethanol solutions over 24 hours, followed by 11 and 12 hours of xylene and Paraplast embedment, respectively.

To maximize RNA integrity, the *Visium HD* assays were performed in accordance with the *10X Genomics* workflow (CG000684, CG0000685) and condensed down to two days. Tissue preparation, including sectioning (5 µm) of the DLPFC, was carried out 3 days prior to the experimental run. Here, microtome, blades, and water baths were initially decontaminated with 80% ethanol and RNase solutions and air dried for 15 minutes. FFPE blocks were trimmed for a minimum of 50 microns prior to collection. Sections were floated for a maximum of 5 minutes at 42°C. Once mounted, sections were left to air-dry for 10 minutes before baking for 3 hours at 42°C. Slides were subsequently stored under a vacuum desiccator until run.

On Day 1, slides were baked at 60°C for 2 hours, then deparaffinized with xylene and rehydrated with graded ethanol solutions. Indirect co-immunofluorescence was subsequently carried out following blocking to visualize amyloid-β (82E1, IBL-Japan, catalog no. IBJP10323E) and phosphor-tau (AH36, StressMarq Biosciences, catalog no. SMC-601). All incubating solutions, including blocking, primary, and secondary antibodies, contained 5% v/v RNase Inhibitor as per the *10X Genomics* protocol. Whole-slide imaging was acquired on the Zeiss Axio Observer Microscope and single-plane Z images were taken. After imaging, slides were decoverslipped and proceeded immediately to probe hybridization. Probe hybridization was enabled for 20 hours rather than the minimum recommended 16 hours.

Day 2 began following probe hybridization with post-hybridization washes, ligation, and post-ligation washes. Here, slides were processed sequentially to maintain thermal consistency during time-sensitive heated washes. Tissues were subsequently maintained on the thermocycler at 4°C during *Visium HD* slide equilibration. For CytAssist-mediated probe release, 15 minutes (instead of the 10 minutes specified by the protocol) were allotted to allow the *Visium HD* capture area to fully dry prior to the run. Subsequent probe extension, probe elution, pre-amplification, and SPRIselect cleanup were performed in direct succession without any pause. Quantitative PCR was then used to determine indexing cycle numbers prior to Sample Index PCR. Samples were amplified, and the resulting indexed libraries were stored at −20°C following a final SPRIselect cleanup. Fragment size and quality assessments were carried out for FFPE libraries prior to sample sequencing using High Sensitivity D5000 TapeStation kit (Agilent Technologies). Libraries were sequenced using a Illumina NovaSeq X Plus using paired-end 150 base pair chemistry (PE150) partial lane sequencing. Each library was expected to generate 150 Gb of data.

### Spatial transcriptomics data preprocessing

#### Samples alignment

Raw FASTQ files in combination with CytAssist eosin images and IF multichannel images were processed by *Space Ranger v3.1.3* with default parameters to obtain count and barcoded location matrices. We used human reference GRCh38-2020-A and *Visium HD* Human Transcriptome Probe Set (v2.0) for alignment. To match coordinate systems between CytAssist and IF images, we used automatic image registration available in the Space Ranger pipeline. In cases where automatic image registration failed, we performed image registration in *Loupe Browser* v8.1.2 using manual landmark selection. For each image, we selected 8-12 matching landmarks with distinct morphological tissue features (for example, tissue borders, folds, and large blood vessels).

#### Nuclei segmentation

A critical step to maximize the interpretation of spatially resolved transcriptomics is the accurate assignment of transcripts to individual cells. Because the *Visium HD* capture area is divided into continuous, fixed 2 µm grids overlaid on the tissue independently of cell boundaries, a single bin can overlap multiple cells, while each cell spans many bins, producing a systematic misalignment between the measured transcriptional signal and the underlying cell positions. To resolve transcripts at the level of individual nuclei, we therefore performed nuclear segmentation on the DAPI image and reassigned the 2 µm bins to the nuclei they overlap.

First, we normalized the DAPI channel images to suppress uneven illumination and background across the field of capture by scaling intensities to the 1-99.2 percentile range with the csbdeep package ^13^. Then, we used the Stardist ^14^ pre-trained fluorescent nuclei model “2D_versatile_fluo” with the wrapper function “model.predict_instances_big” for large images, splitting them into overlapping image blocks (block_size=4096, min_overlap=128, context=128). We used the optimized model parameters recommended by *10X Genomics* guidelines to enable less-confident predictions, with a more conservative overlap threshold (prob_thresh=0.01, nms_thresh=0.001). All segmentation computations were performed on a high-performance cluster node with a GPU, with an average execution time = 2m 20s, depending on microscopy image resolution. Finally, to merge subcellular 2 µm bins into nuclei, we used bin2cell ^15^ to assign bins to their corresponding nuclei and aggregate them at single-cell resolution. Before assignment, we removed non-expressing bins (at least 1 UMI) and genes with expression detected in a low number of bins (expressed in < 3 cells). Then, bins were assigned to corresponding Stardist nuclei labels using the “insert_labels” function and expanded to detect cytoplasmic transcriptomic signals with “expand_labels” function, with a default expansion level of 2 bins. We omitted secondary label annotations obtained from transcriptomic expression, as these labels appeared excessive and noisy. The merged bin2cell output dataset contains 796,374 cells.

#### Quality control

Quality control and processing were performed using the *scanpy* package ^16^ for FFPE and FF sample batches from the frontal cortex region. The numbers of UMIs and genes, and the percentages of mitochondrial and ribosomal content were obtained using “pp.calculate_qc_metrics” (qc_vars=[“mt”, “ribo”], log1p=True). Additionally, we used the bin2cell calculation to estimate the number of 2 µm bins per cell as a proxy for cell size. To determine optimal quality metric levels, we applied quantile-based thresholds for individual samples. For the number of UMIs and genes, we set the QC threshold at the 10th percentile (low-quality cells); for the number of bins between the 5th and 95th percentiles (segmentation errors, background noise, or large artifacts). A fixed threshold of 10% was set for the mitochondrial content percentage. Our initial dataset after QC filtering included 578,159 total number of cells. Sample-specific QC threshold values are available in Supplementary Table 3.

#### Data integration and annotation

We performed data integration using highly variable genes (HVGs) identified from a reference single-nucleus RNA-sequencing dataset, rather than recalculating HVGs across all samples in the merged dataset. This strategy was designed to minimize sample-to-sample heterogeneity and facilitate a robust clustering and annotation of the major, well-represented brain cell populations.

To define the HVG set for spatial transcriptomics integration, we used an snRNA-seq reference dataset (AUSBB cohort) ^17^ in which eight major cell types had been confidently annotated (excitatory neurons, inhibitory neurons, microglia, astrocytes, oligodendrocytes, oligodendrocyte precursor cells, endothelial cells, and vascular leptomeningeal cells). For each cell type, we selected the top 250 marker genes using the FindAllMarkers function in Seurat (v4.4.0) ^18^. The union of these lists yielded 1,612 unique genes, which we used as the reference HVG set (Supplementary Table 5). We excluded *CLU*, *MBP*, *GFAP* from HVGs as the probes used to capture the expression of these did not resemble single-cell cell-type distributions. The reference HVGs that were expressed in fewer than 10 cells were also excluded. Next, we applied additional QC filtering for the number of UMIs and excluded cells that did not express any HVGs. The final merged spatial dataset contained 1,548 genes and 558,145 cells. To reduce technical bias between samples and preservation methods, we performed scVI integration using the respective sample ID annotations for batch correction. The scVI model was trained (‘model.train’) on a GPU-enabled HPC node for 500 epochs with default early stopping. 10% of the dataset was used for model validation. We used scVI embeddings to compute nearest neighbors and a low-dimensional UMAP embedding and calculated unsupervised Leiden clusters on a log-normalized dataset with default parameters for scanpy functions. Then, to remove remaining low-quality clusters, we performed iterative clustering at a low resolution (res= 0.1) and excluded clusters containing fewer than 100 cells that failed to exhibit coherent gene expression profiles or spatial enrichment within a defined anatomical region. For each iteration, nearest neighbors, unsupervised clusters and UMAP reductions were recalculated. The final cleaned integrated dataset contains 530,172 cells annotated into 12 clusters (res=0.7), for which we identified differentially expressed marker genes using sc.tl.rank_genes_groups function with the following parameters: groupby=“scvi_leiden_0.7”, method=’wilcoxon’, use_raw=False. Clusters were then annotated by cell type using selected markers (neurons - *SYT1*, *PVALB*, *GAD1*, *GAD2*, *RTN3*, *SST*; astrocytes - *AQP4*; oligodendrocytes - *PLP1*, *MOG*; microglia - *C3*, *CD74*, *SPP1*, *C1QB*, *S100A9*, *S100A8*; vascular and endothelial cells - *TAGLN*, *CLDN5*, *MYL9*, *EPAS1*). The necessity of an iterative cleaning process was driven by differences in QC metrics across preservation methods (Supplementary Table 4). Cumulatively, 47,528 cells (mainly from sample ffpeAD3_S1) were removed in FFPE samples compared to 459 cells in FF samples (18.6% and 0.1%, respectively).

### Determining the optimal sequencing depth for human brain tissues

We performed saturation analysis to identify the most cost-effective number of reads to sequence the FF brain tissues that maximize transcriptional profiling. To simulate how changes in sequencing depth would affect the quality of transcriptomics data, we used a subsampling approach to perform saturation analysis ^19^, using the two slides (S1 and S2) we generated from donor AD2. Original sequencing was performed with a target level of 600 million reads. Then, for both samples, we performed additional sequencing of 700 million reads using the aliquots of the same libraries sequenced originally, yielding ∼1.2 billion reads for the final objects. Subsampling was performed with the *seqtk* tool (GitHub, https://github.com/lh3/seqtk) by selecting 10-90% random reads from the total FASTQs with 10% increase steps. To keep the matching read rows, we used the same random seed (“-s 123”) for both R1 and R2 files. For each subsampled FASTQ file, we ran *Space Ranger* with the same parameters as for the full objects to obtain subsampled count matrices. QC metrics were calculated with *scanpy* function “pp.calculate_qc_metrics” per 2, 8,16 µm bins and nuclei resolutions. In parallel, we were leveraging transcriptomics for cellular segmentation. Thus we used the same DAPI-based nuclei annotations for all sequencing depth estimates. Sequencing saturation was calculated with the following formula: 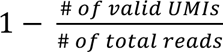.

### Ultima Genomics UG 100 sequencing

*Library conversion.* To create libraries compatible with the *Ultima Genomics UG 100* sequencer, original libraries (25 µL per sample) with Illumina adapters were obtained from an optimized *10X Genomics Visium HD* protocol and subjected to a single round of indexing PCR amplification (*Ultima* protocol D1001055). The reaction was prepared using a compatible PCR master mix containing the xGen UG-tR1 Indexing Primer v2, which served as the forward primer, and the xGen UG-srR2 Universal Primer v2, which served as the reverse primer. Following thermocycling, the amplified libraries underwent a magnetic bead clean-up using AMPure XP beads at a 1.2x volumetric ratio to ensure purity. The beads were separated on a magnet, washed twice with 80% ethanol, and allowed to air-dry. The final UG-compatible libraries were then eluted in 150 µL of buffer (10 mM Tris, pH 8.0). Before sequencing, library quality and quantity were verified by measuring DNA concentration with a Qubit fluorometer and average base pair length with a fragment analysis instrument. Final libraries were sequenced on the Ultima Genomics UG 100 platform using 1/3 of full wafer.

*Sequencing data processing.* By default, the *Ultima Genomics UG 100* sequencer outputs CRAM file containing 300 bp single-end reads. To convert the single-read data generated into a paired-end FASTQ format compatible with the *10X Genomics Space Ranger* alignment pipeline, on-instrument data processing was executed using the *Ultima Genomics Trimmer* software (*Ultima* protocol D1001122). During this process, the algorithm removed the *Ultima* adapter sequences, any conserved sequences remaining from the conversion protocol, and application-specific regions. Simultaneously, the software identified and isolated the *10X Visium HD* spatial barcodes and the Unique Molecular Identifiers (UMIs) while demultiplexing the samples using the *Ultima* barcode sequence. To simulate paired-end reads, Read 1 (R1) was constructed to contain the extracted spatial barcode along with the UMI, while Read 2 (R2) was constructed to contain the corresponding ligated probe sequence. Converted FASTQ files were processed with the Space Ranger v3.1.3 downstream pipeline as described previously.

To generate sequencing depth-matched *Ultima* FASTQ files, *Space Ranger*– aligned BAM files were downsampled using *Samtools* 1.24 ^20^. Target number of reads was calculated as the proportion of reads using the following formula: 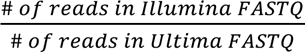. A random seed was provided for reproducibility (seed = 123) and to ensure matching read rows. Downsampled BAM files were converted to FASTQs using the *bamtofastq* tool (from the *Space Ranger* toolkit). The resulting FASTQ files were processed with *Space Ranger v3.1.3* using the same parameters as the original files.

## Results

### *Visium HD* protocol optimization for fresh-frozen human post-mortem brain

To improve preservation of FF post-mortem brain tissue during cryosectioning, tissue fragments from post-mortem brains that had been directly frozen at −80 °C after autopsy were embedded in OCT medium. First, to determine whether OCT embedding affected RNA integrity, we dissected frontal cortex fragments from two donors diagnosed with AD and divided each fragment into two smaller pieces: one used immediately for total RNA isolation, and the other embedded in OCT, as described in the Methods section. Approximately four cryosections (25 µm each) of the OCT-embedded tissue were collected for RNA extraction, and RNA quality from both conditions was assessed using the TapeStation system (Figure 1A). Of the two analyzed samples, the RINe score decreased after OCT embedding in one donor (AD2), from 6.6 to 5.3. Although RNA from donor AD1 did not show substantial changes in RINe after OCT embedding, its total RNA was overall more degraded relative to AD2, with a RINe of 4.2 (Figure 1B). This suggests that OCT embedding may great a negative effect on higher-quality samples.

**Figure 1:**
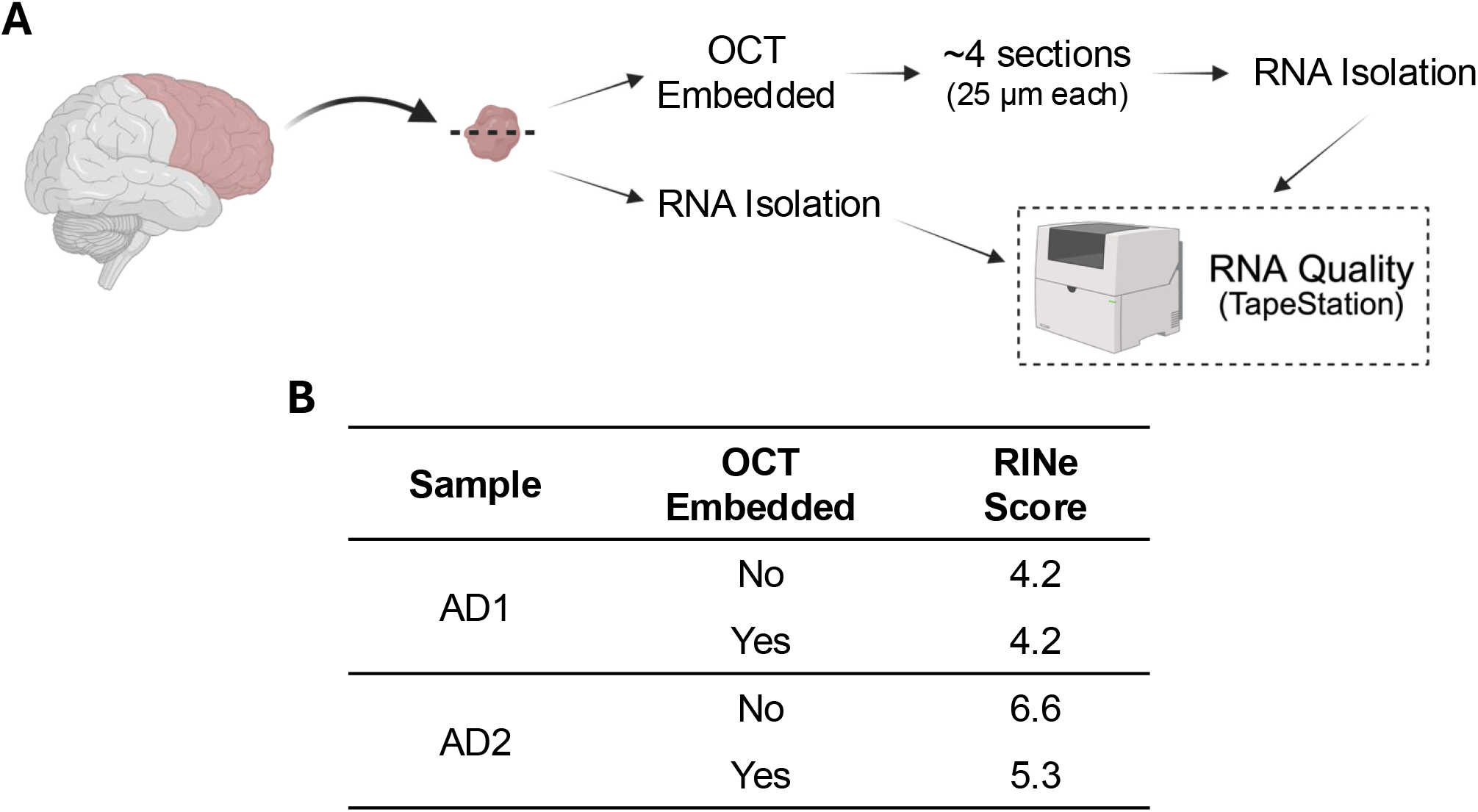
RNA quality assessment of human post-mortem brain tissue. (A) A tissue fragment collected from fresh-frozen (FF) dorsolateral prefrontal cortex (DLPFC) was divided into two smaller fragments. One fragment was embedded in optimal cutting temperature (OCT) compound prior to cryosectioning (∼4 sections of 25 µm each) and subsequent RNA isolation, while the second fragment was directly used for RNA extraction. RNA integrity from both conditions was assessed using the TapeStation system. (B) RNA Integrity Number equivalent (RINe) values for two FF samples before and after OCT embedding. Figure creaded with *BioRender.com*.

Because the *10x Genomics Visium HD* workflow includes several recommended stop points that may prolong tissue handling and storage, we next evaluated whether these pauses affect spatial transcriptomic library quality in FF post-mortem human brain tissue. To do so, we compared the standard workflow containing the recommended stop points with a modified continuous workflow performed without interruptions (Figure 2).

**Figure 2:**
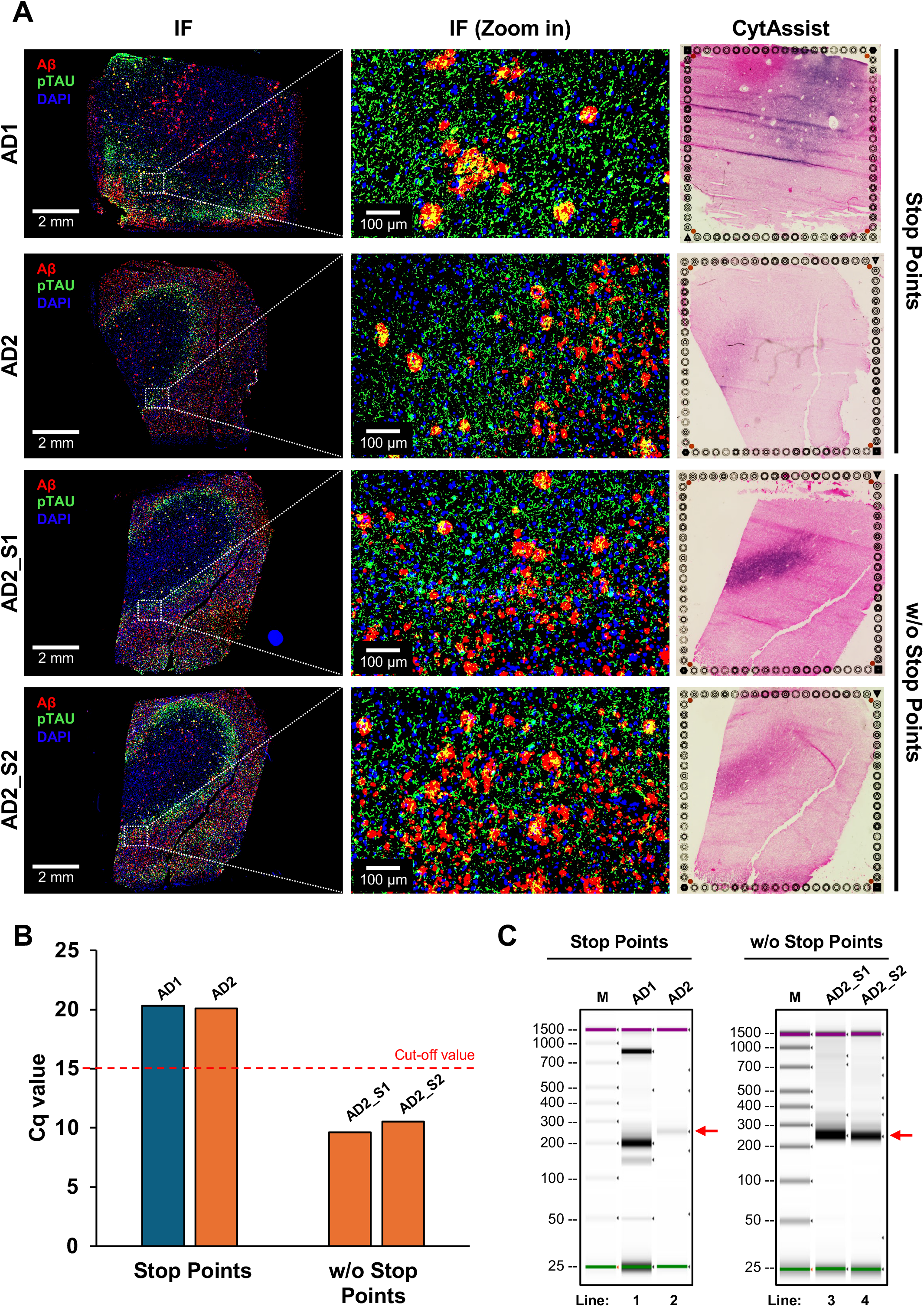
*Visium HD* protocol optimization for human post-mortem brain tissue. (A) Immunofluorescence images from two Alzheimer’s disease samples (AD1 and AD2) stained for amyloid-β (Aβ), phosphorylated tau (pTau), and DAPI, alongside their corresponding eosin-stained images generated by CytAssist, following the original *Visium HD* protocol, including the use of the recommended stop point between staining and coverslip removal. Two additional adjacent sections from AD2 (AD2_S1 and AD2_S2) were stained for the same targets and processed using the optimized workflow without any stop points; corresponding eosin-stained images are shown in the last column. (B) Cq values obtained from qPCR quantification of the pre-amplified cDNA for samples processed using the recommended stop points and 16-hour probe hybridization (left), compared to samples processed without stop points and with extended probe hybridization (20 hours; right). (C) Final library quality assessment by TapeStation for samples processed using the original *Visium HD* protocol (left panel) and the optimized protocol (right panel). The expected library fragment size (∼250 bp) is indicated by red arrows.

As reference condition, FF post-mortem brain tissue sections from two AD donors (AD1 and AD2), were processed according to the standard *10X Genomics Visium HD* user guide. After cryosectioning, the mounted slides were stored at −80 °C for one week before immunofluorescence staining. Tissue sections from both donors were stained for Aβ and pTau (1-hour incubation with primary antibodies and 20 minutes with secondary antibodies, as recommended by the *10X Genomics* protocol), along with DAPI, and were imaged by fluorescence microscopy. Prior to coverslip removal for probe hybridization, the stained slides were stored flat in the dark at 4 °C for 48 hours, corresponding to one of the stop points recommended in the user guide. The IF staining for both AD samples showed the expected AD-associated pattern, with Aβ plaques and neurofibrillary tangles (NFTs) distributed throughout the tissue, with higher density in the gray matter (Figure 2A).

After coverslip removal, the probes were hybridized for 16 hours, following the user guide recommendations, and the protocol was continued until the post-sample index PCR cleanup (SPRIselect) step without additional stop points. However, despite successful visualization of tissue pathology, downstream library quality metrics indicated suboptimal assay performance. The cycle number determination by qPCR, which can be used as an indicator of probe capture efficiency, resulted in quantification cycle value (Cq) of 20.3 and 20.0 for AD1 and AD2, respectively (Figure 3B, left bars), substantially exceeding the recommended threshold of ≤15 cycles. The post-library QC for AD1 showed a major band at ∼200 bp and several fainter bands at ∼150 bp and ∼50 bp (Figure 2C, Line 1, left panel). The library QC for AD2 yielded a single faint band near the expected fragment size of at ∼250 bp (Figure 2C, Line 2, left panel). Together, these findings suggested that one or more steps in the standard workflow, including the recommended stop points, may compromise library generation from FF post-mortem brain tissue.

**Figure 3:**
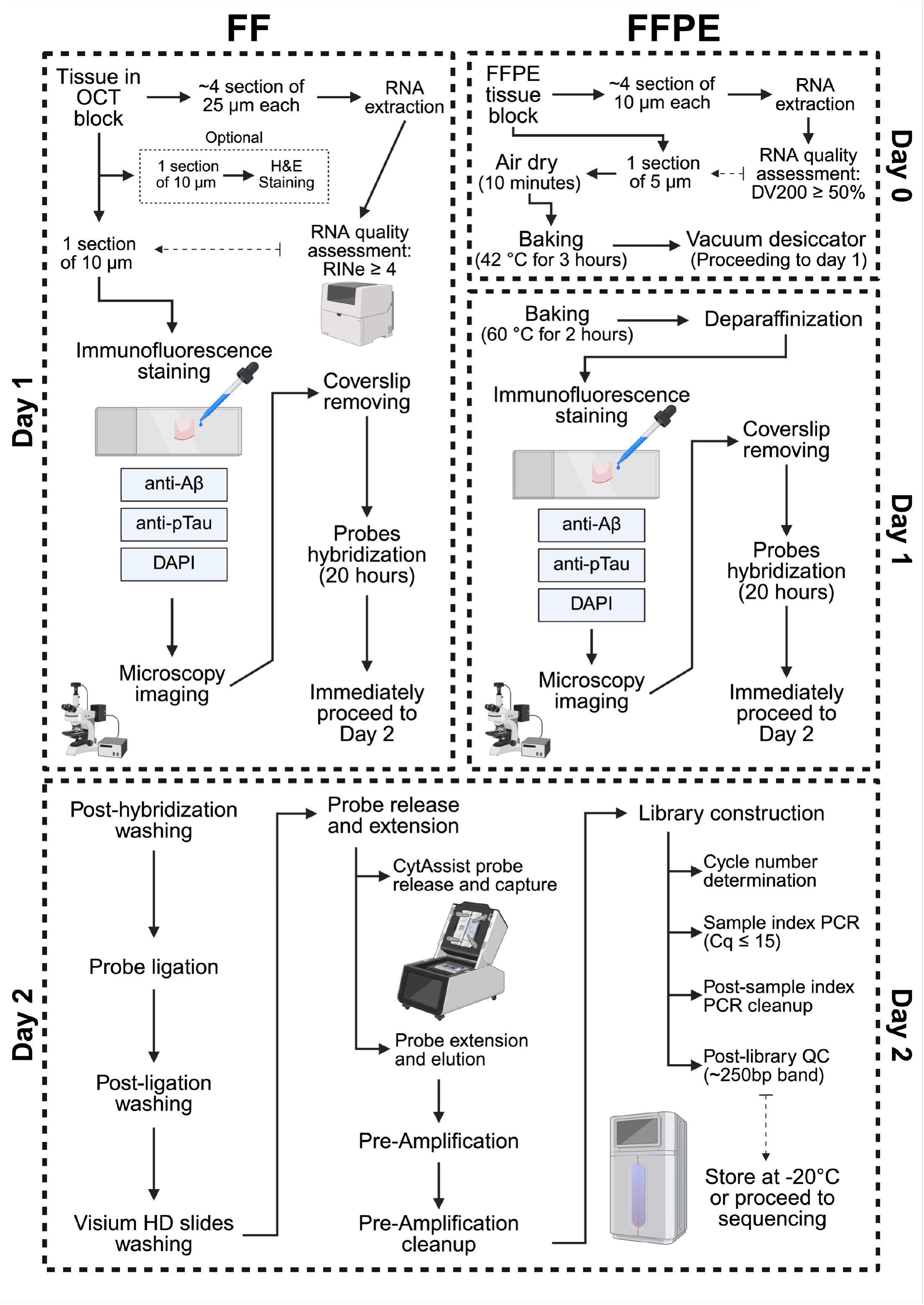
Optimized workflow for fresh-frozen (FF) and formalin-fixed paraffin-embedded (FFPE) human brain tissue processing for spatial transcriptomics using the *Visium HD* platform. Figure creaded with *BioRender.com*.

To directly test this possibility, we next processed tissue using a modified workflow performed continuously over two consecutive days without any of the recommended stop points. Because AD2 exhibited the highest RNA quality following OCT embedding (Figure 1B), two consecutive tissue sections from this donor (AD2_S1 and AD2_S2) were selected for protocol optimization. In addition to eliminating stop points, probe hybridization time was extended from 16 to 20 hours to maximize transcript capture efficiency (Figure 2).

After applying these modifications, we observed comparable AD IF staining patterns in the two consecutive sections and consistent eosin staining for CytAssist (Figure 2A, rows 3 and 4). In contrast to the standard workflow, library QC for the samples prepared under the optimized workflow showed marked improvements. The Cq values decreased to 9.6 and 10.5 cycles for AD2_S1 and AD2_S2, respectively (Figure 2B, right bars). The post-library QC analysis also improved substantially, with both consecutive slices showing a prominent band at the expected fragment size of ∼250 bp (Figure 2C, Lines 3 and 4, red arrows in the right panel).

Due to the success of the *Visium HD* protocol optimization, three additional FF human post-mortem brain samples (AD3_S1, AD4_S1, and AD5_S1) were selected and processed according to the optimized workflow (Figure 3). All three tissue sections were stained for Aβ and pTau, in addition to DAPI, and together with the eosin staining obtained from CytAssist, demonstrated good overall tissue quality (Figure 4A). Correlation analysis between RNA quality (RINe) and the Cq value of the pre-amplified product showed a strong relationship between RNA integrity and amplification efficiency (Figure 4B). Specifically, cDNA generated from lower-quality RNA samples (lower RINe) exhibited higher Cq values, indicating the need for a greater number of PCR cycles to generate the final sequencing libraries with the expected size (∼250 bp) (Figure 4C).

**Figure 4:**
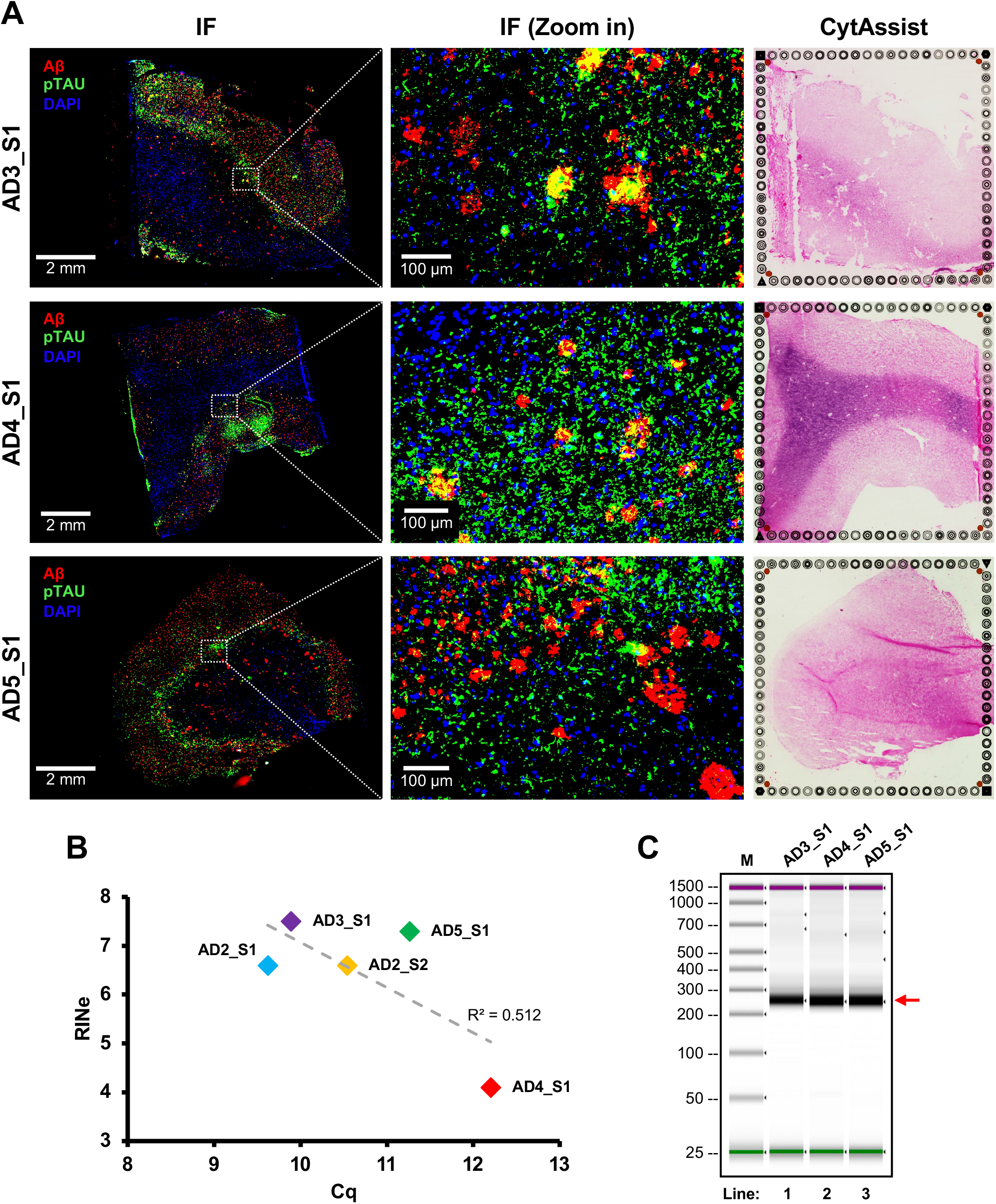
RNA quality (RINe) and cDNA amplification (Cq) correlate with final library quality. (A) Immunofluorescence images from three additional Alzheimer’s disease samples (AD3_S1, AD4_S1, and AD5_S1) stained for amyloid-β (Aβ), phosphorylated tau (pTau), and DAPI, alongside their corresponding eosin-stained images generated by CytAssist, following the optimized *Visium HD* protocol. (B) Correlation between RINe score and cDNA pre-amplification Cq values obtained for each fresh-frozen sample, indicating a direct relationship between RNA quality and the number of PCR cycles required for final library amplification. (C) Final library quality assessment by TapeStation for samples processed using the optimized *Visium HD* protocol. The expected library fragment size (∼250 bp) is indicated by red arrows.

Direct comparison of the two workflows demonstrated that the modified continuous protocol consistently outperformed the standard workflow. The optimized protocol produced substantially lower Cq values and generated libraries with fragment size distributions centered around the expected ∼250 bp peak, whereas the standard workflow yielded elevated Cq values and poor-quality libraries. Together, these results indicate that the workflow modifications, most notably the elimination of protocol stop points, together with extended probe hybridization, substantially improve library construction from FF post-mortem human brain tissue and provide a more robust approach for Visium HD spatial transcriptomics.

### Finding minimal and optimal sequencing depth

To assess *Visium HD* data quality before downstream processing, we performed a sequencing-depth simulation, followed by saturation and quality-control analyses, on FF brain samples. We conducted analysis using two consecutive tissue sections from the same donor (AD2_S1 and AD2_S2) to minimize sample-specific bias. Consistent with the increased susceptibility of post-mortem human FF brain tissue to RNA degradation, these samples provided a stringent test scenario for the sequencing depth effect. Assessment of tissue coverage revealed that 76.1% of the *Visium HD* capture area was occupied by tissue in AD2_S1, whereas AD2_S2 showed coverage of 82.8% of the capture area. We sequenced libraries to target 562.5 million reads per sample, yielding 548 and 533 million reads in the AD2_S1 and AD2_S2 samples, respectively. This depth was close to the recommended target sequencing depths for the libraries, which were expected to be 532.7 and 579.6 million for AD2_S1 and AD2_S2, respectively, according to the *Visium HD* Spatial Gene Expression Reagent Kits User Guide (CG000685, Rev. D; “Sequencing Depth” section). While both samples met the requirements for minimal sequencing depth, we decided to assess whether higher sequencing depth could significantly affect data quality. We followed up with the next round of sequencing of the same libraries, yielding an additional 625 million reads per sample. The final number of sequences totaled 1,210.8 and 1,240.4 million reads for AD2_S1 and AD2_S2, respectively. To modulate intermediate sequencing depths, we performed a simulation of sequencing with lower depth by subsampling reads from FASTQ files (10-90% of the total reads with 10% increments) for each sample before and after additional library sequencing.

Sequencing saturation analysis (Figure 7A) demonstrates that sequencing levels recommended by 10x guidelines reached 81% and 88% saturation for the AD2_S1 and AD2_S2 samples, respectively. Doubling the sequencing depth increased sequencing saturation by 9.2% (AD2_S1) and 6.8% (AD2_S2), suggesting that both samples had reached the point where additional sequencing would not recover much new information. By quantifying subsampled data, we were able to analyze the effects of sequencing depths on transcriptomics data quality metrics, measured by the number of UMIs and the number of genes per bin or nuclei (Figure S1). Simulation results demonstrate that the inflection point in the plots of mean number of UMIs and genes was reached at a significantly lower sequencing depth (260-270 million reads) compared with sequencing saturation for both samples, independent of their aggregation levels. We obtained similar results when analyzing the total number of expressed genes (Figure 7B). However, we noticed that switching from spatial bins to the individual-nucleus transcriptomics context reduced the average number of expressed genes by 1.8% (294 genes). Gene set enrichment analysis (GSEA) showed that genes are involved in multiple processes associated with plasma membrane and neurotransmitters activity – signal transduction (OR1A2; OR4E2; OR14J1; CALML5), neuroactive ligand-receptor interaction (GHSR; EDN2; TAAR9; TAAR6; RXFP4), cytokine-cytokine receptor interaction (IL1RN; CCL23; CXCL9; IL2; CCL21), and GPCR ligand binding (CXCL9; TAS2R42; CCL3L1; RXFP4; HTR2B). This indicates that these genes could be used to increase the accuracy of nucleus boundary prediction, although their expression levels are low, which significantly reduces their applicability.

Overall saturation analysis demonstrates that the minimal sequencing depth guidelines provided by *10X Genomics* result in high sequencing saturation (>80%) in fresh-frozen human postmortem brain tissue. At the same time, our analysis of transcriptomics data processing QC metrics shows we achieved appropriate levels with a significantly lower sequencing depth (∼53% lower) than the recommendations, assuming an initial high RIN score. We also demonstrated that, for selected tissues, increasing sequencing depth did not affect sequencing saturation or data QC metrics, suggesting that samples had reached saturation and no additional gene expression was detected.

### *Visium HD* data processing

Although the *10x Genomics* Space Ranger pipeline provides a framework for primary processing of *Visium HD* data, several challenges remain for the analysis of post-mortem human brain tissue. First, while Space Ranger implements image-based nuclei segmentation using DAPI staining, the default workflow does not extend segmentation boundaries to capture nuclei-adjacent cytoplasmic transcripts, potentially reducing transcript recovery and sensitivity for cell-type identification. Second, substantial variability in RNA quality, transcript yield, and tissue preservation can lead to large differences in data quality across samples, particularly when integrating FF and FFPE tissues. Finally, standard highly variable gene (HVG)-based integration approaches may perform poorly in spatial transcriptomic datasets because technical and sample-specific biases often dominate the set of detected HVGs, obscuring shared biological signals. To address these challenges, we developed a unified pipeline for the initial QC and analysis of whole-transcriptome *Visium HD* data that is independent of tissue preservation method and integrates nuclei segmentation with cytoplasmic expansion, adaptive quality control, reference-guided sample integration, and cell-type annotation within a single workflow (Figure 7C). Beginning with raw Space Ranger count data, the pipeline generates an integrated Scanpy object suitable for quality assessment and downstream analyses.To evaluate the robustness of our approach, we expanded our dataset with three additional fresh-frozen (Figure 4) and two FFPE (Figure 5) post-mortem human brain tissues. The final cohort comprises seven libraries originating from DLPFC samples (5 FF and 2 FFPE). All donors were diagnosed with AD, and the analyzed tissues exhibited substantial levels of AD-associated pathologies (Supplementary Table 1).

**Figure 5:**
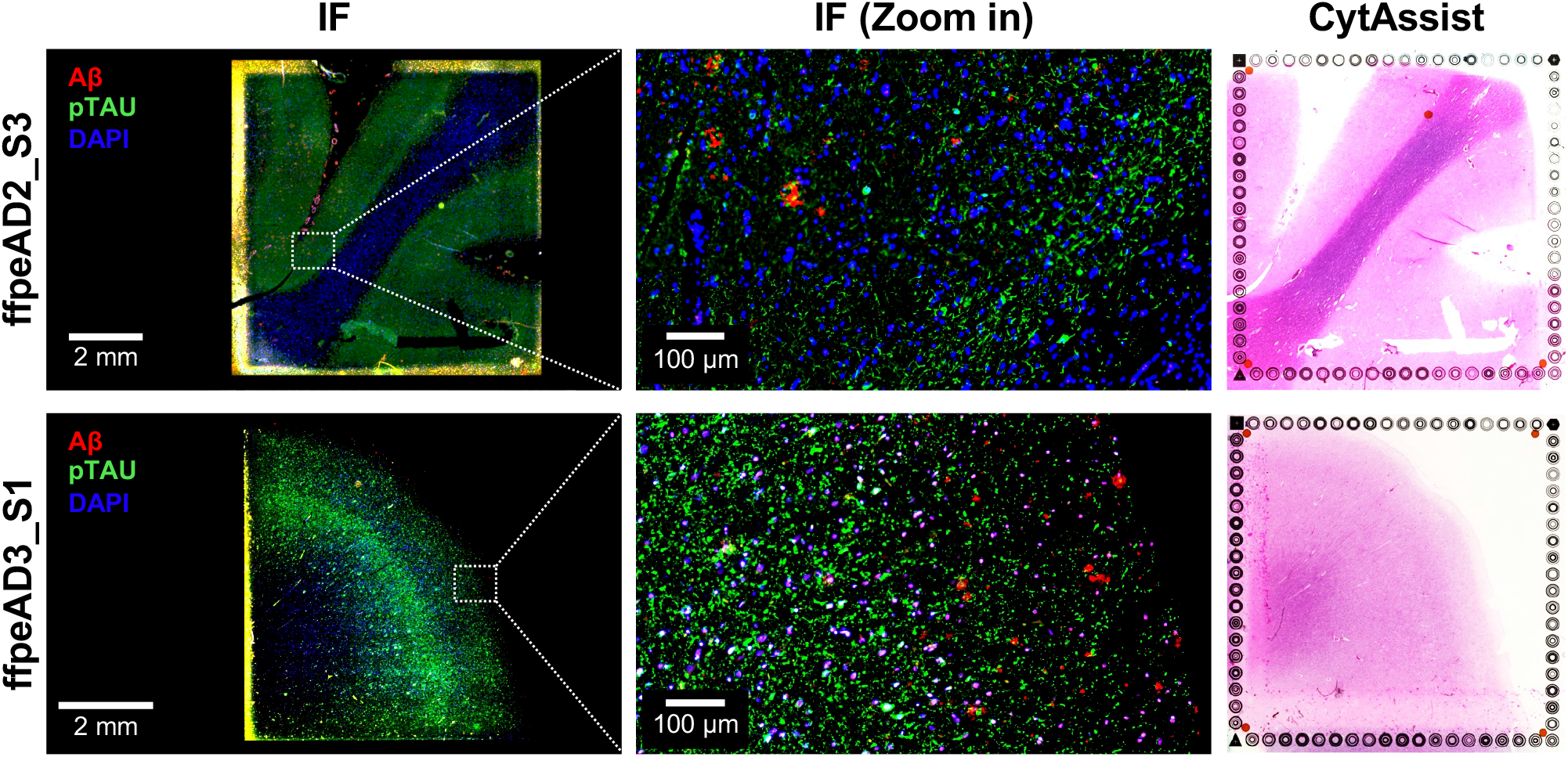
Immunofluorescence staining of FFPE human post-mortem brain tissue. Immunofluorescence images from two Alzheimer’s disease samples (FFPE_AD2_S3 and FFPE_AD3_S1) stained for amyloid-β (Aβ), phosphorylated tau (pTau), and DAPI, alongside their corresponding eosin-stained images generated by CytAssist.

From the default Space Ranger quantification procedure for *Visium HD* spatial sequencing data, we obtained gene transcript counts across a continuous tissue grid subdivided into smaller barcoded segments (bins) with multiple resolutions (2, 8, and 16 µm). However, such bins are not explicitly associated with individual cells and may capture empty extracellular space or only partial signals from single cells and cell aggregates (doublets). To address this limitation, we reasoned that nuclei-adjacent bins would more reliably capture cell-type-specific transcriptomes, and we performed nuclear segmentation based on DAPI staining and applied a 4-µm nuclear border expansion to capture nuclei-adjacent cytoplasmic signals. We then compared quality metrics of the resulting cell aggregates to those from the bin-based resolutions produced by the Space Ranger pipeline. We found that 8-µm spatial bins best approximate the distributions of the nuclei-adjacent cytoplasm data (Figure 7D and S2A). Although median and interquartile range (IQR) values were similar across both approaches, cell aggregates exhibit unimodal distributions of unique molecular identifiers (UMIs) and detected genes, particularly at the lower counts, reflecting the exclusion of empty areas. We observed no differences in the percentage of mitochondrial transcripts.

**Figure 6:**
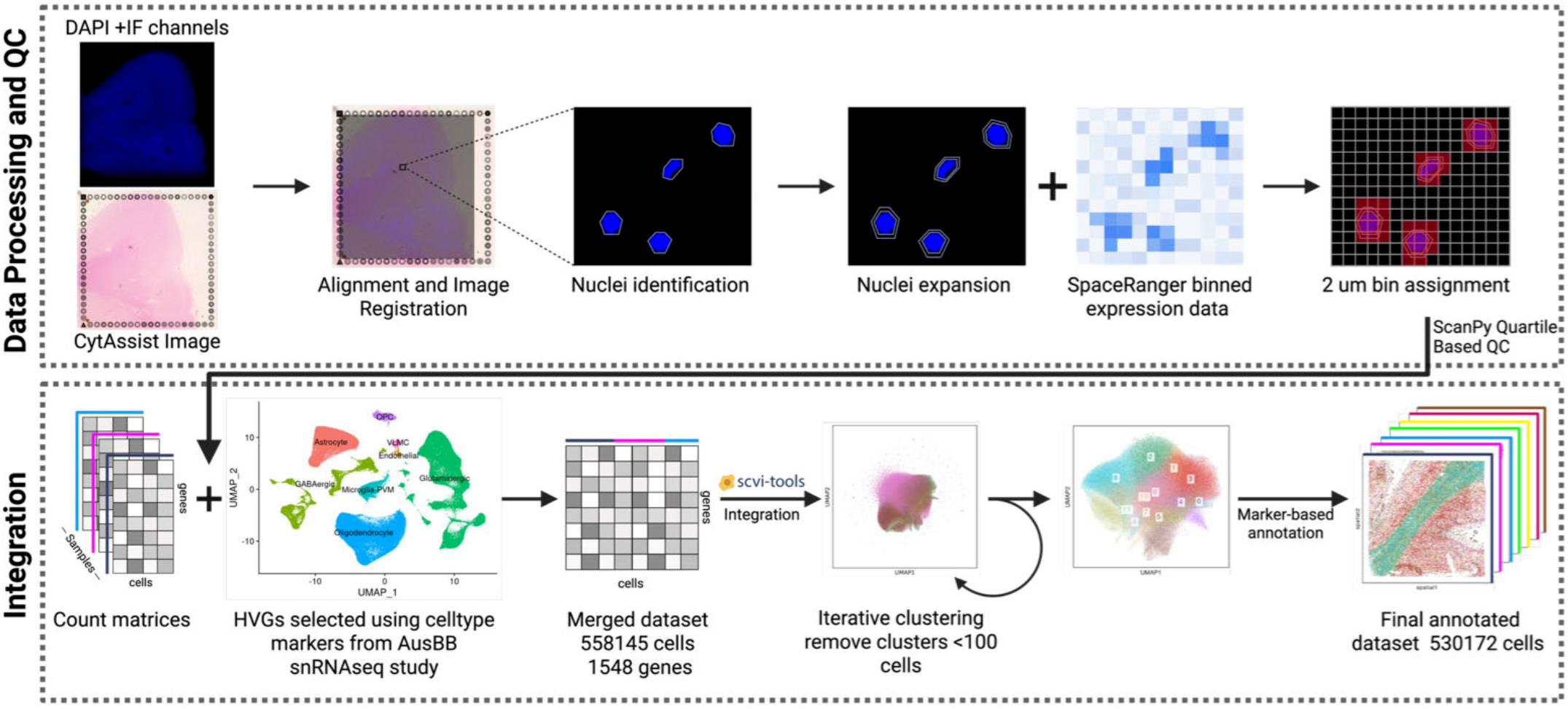
Overview of the *Visium HD* data processing pipeline. The immunofluorescence channels, including DAPI, were aligned to the CytAssist image to register them to the *Visium HD* capture area using Loupe Browser. Nuclei identification was performed using StarDist on the DAPI IF channel. Bin2cell was used for nuclear expansion and to assign 2 µm binned Space Ranger outputs to expanded nuclei resulting in single cell count matrices for each sample. Sample-wise QC was performed using scanpy prior to integration across samples. When integrating, HVGs were selected using cell type markers identified in the AUSBB snRNA-seq dataset. Integration was performed using scVI. Following integration, cells were clustered iteratively, removing any clusters with fewer than 100 cells prior to reclustering. The resulting annotated dataset contains 530,172 cells across 12 clusters. Representations of nuclei detection and bin assignment are schematic, drawn on a magnified DAPI region from a representative slide.

**Figure 7:**
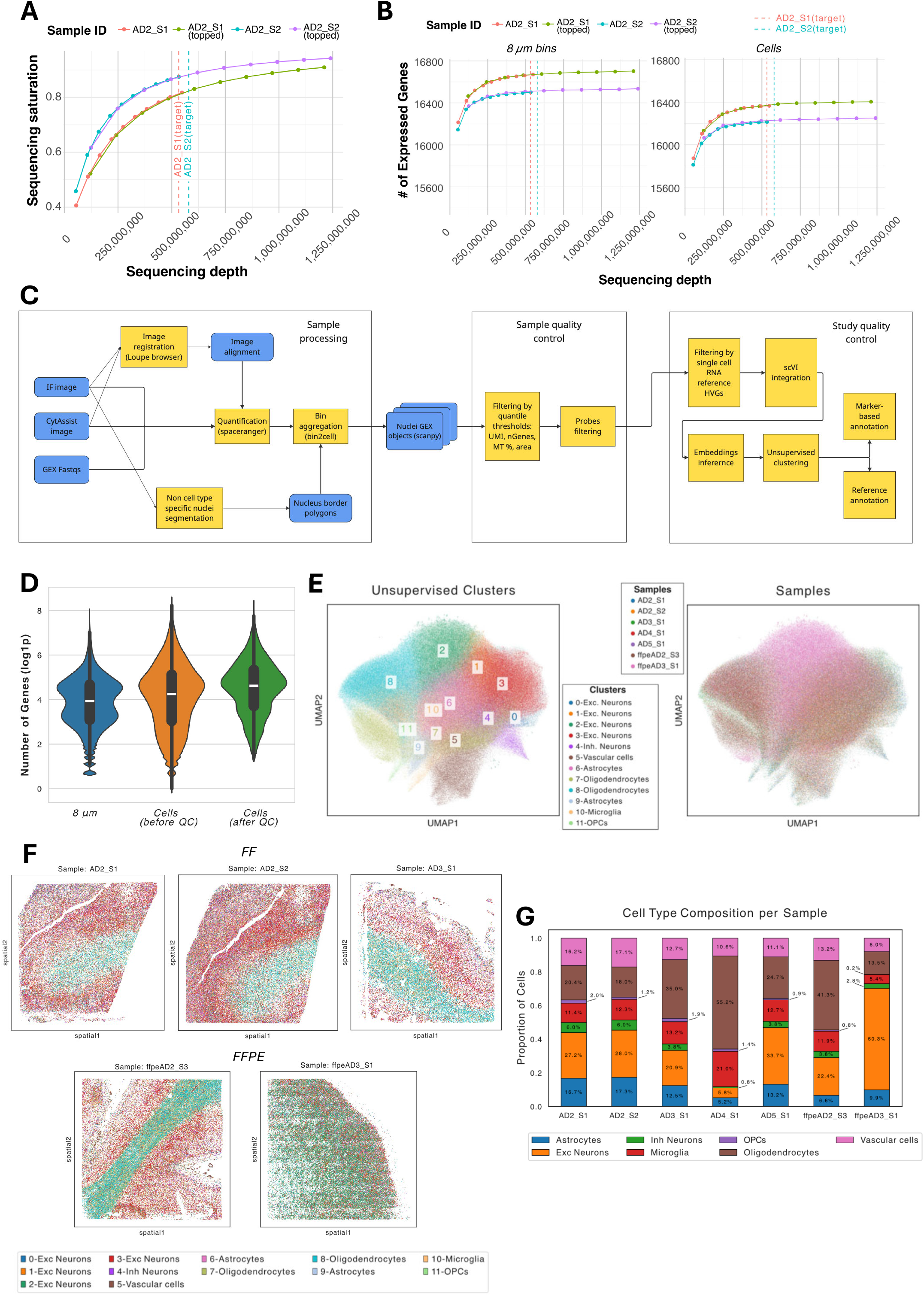
Data processing and quality control for the *Visium HD* human brain dataset. (A) Sequencing depth association with sequencing saturation for fresh-frozen brain tissues. Intermediate metrics were simulated by subsampling FASTQ files and quantifying them using the Space Ranger pipeline. (B) Association between sequencing depth and the number of total expressed genes for 8 µm bins and segmented cells. For (A) and (B), the target levels represent the 10X minimal recommendations for sequencing depth. (C) Flow diagram for data processing and QC pipeline. (D) Violin plots comparing distributions of the number of genes between different aggregates – 8 µm bins and segmented cells. Pre-QC filtering segmented cells distribution has a uniform shape in the lower band compared to the 8 µm bins. QC filtering for segmented cells highlights minimal effect on overall distribution shape. The number of genes was scaled using the ln (X+1) transformation. (E) Integrated UMAP plot with identified 12 unsupervised clusters with cell type annotations and UMAP with sample distribution. The final dataset contains a total of 530,172 cells. Cluster 2 is primarily associated with the ffpeAD3_S1 sample. (F) Spatial plots with annotated cells illustrating tissue architecture and highlighting the distinction between gray and white matter areas. (G) Composition of major cell types’ proportions per sample. Replicate samples AD2_S1 and AD2_S2 (from the same tissue block) show similar cell-type proportions.

We noted divergent QC metrics across samples (Supplementary Table 2) and used a derived quantile-based threshold approach to filter individual samples by the number of UMIs, genes, and cells (2 µm bins per cell). This strategy prevented the removal of a substantial number of cells from the general lower-quality FFPE sample ffpeAD3_S1. In contrast, as median mitochondrial content per sample was low (≤ 5%) and showed no effect on downstream analysis, we used a fixed threshold (10%) to identify tissue areas damaged during preservation or sample preparation. Consequently, we removed 36,382 cells (49.7%) due to high mitochondrial content from sample AD4_S1, in which we observed damaged gray matter area that may have originated from the initial freezing of the tissue, whereas most cells in the white matter showed no damage, and passed QC (Supplementary Table 3).

Although all samples were obtained from the same brain region, we anticipated substantial batch effects due to assay-specific technical biases and differences in brain bank-specific tissue-handling procedures. To estimate transcriptional similarity across samples, we first identified highly variable genes (HVGs) within each sample and then quantified pairwise similarity using the Jaccard index (Figure S2B). The resulting heatmap revealed minimal overlap in HVGs between samples, even for those libraries derived from the same donor. These observations indicate that downstream analyses of such data, even with batch correction, are prone to yield noisy principal components and poorly resolved clusters that primarily reflect sample-specific probe bias rather than the underlying biology.

To enable integrated analysis for cell-type annotation, we implemented a reference-based HVGs selection strategy in conjunction with training of the scVI model embeddings. We derived the reference gene set from a single-nucleus RNA-seq dataset of post-mortem parietal lobe tissue (71 individuals) from the AUSBB cohort, comprising patients with late-onset AD (LOAD), autosomal dominant AD (ADAD) and age-matched controls. Because our primary objective was cross-sample integration to facilitate cell type annotation, we defined reference HVGs as the top 250 differentially expressed (DE) marker genes associated with the major cell type clusters in the reference dataset (Supplementary Table 5). Although the resulting reference gene list showed limited overlap with HVGs identified in the spatial transcriptomics samples, integration based on this reference substantially reduced inter-sample heterogeneity and yielded coherent, shared unsupervised clusters across samples (Figure 7E).

We next evaluated whether this integration strategy effectively mitigated batch effects. For each sample, we computed cell density distributions in the integrated UMAP embedding space (Figure S2C). Apart from sample ffpeAD3_S1, UMAP density plots showed a fair distribution of cell types among samples. The final integrated dataset comprised 530,172 cells and contained 12 clusters. Cluster identities were assigned to major cell types based on canonical marker genes, differential expression analysis, and pathway enrichment analysis (Figure S3A-C). These complementary validation approaches were concordant and supported the robustness of the inferred cell type annotations.

Spatial plots with cell-type annotations (Figure 7F and S4A) revealed distinct morphological features across tissues, highlighting both unique and shared characteristics. Tissues contained a clear demarcation between gray and white matter regions, with the white matter exhibiting an abundance of oligodendrocyte cell clusters. The gray matter predominantly contains neuronal subpopulations, with distinct clusters (Clusters 0 and 1) reflecting transcriptional differences between upper and deeper cortical layers. In tissues with distinct large blood vessels, such as sample ffpeAD2_S3, we observed well-defined vascular structures, primarily composed of cells from Cluster 5, which expressed vascular and endothelial cell markers. Cell type compositional analysis (Figure 7G) demonstrated cellular heterogeneity across samples, thereby validating the observed spatial tissue architecture. A comparison of serial slices from the same tissue block (AD2_S1 and AD2_S2) showed no significant differences in cell populations, consistent with the defined clusters capturing genuine biological signatures rather than sample-specific or preservation artifacts. Notably, sample AD4_S1 exhibited an increased proportion of oligodendrocytes (55.2%), which we hypothesize reflected specific gray matter damage. Conversely, sample ffpeAD3_S1 displayed a high abundance of excitatory neurons (60.3%), predominantly belonging to Cluster 2, which is largely absent in other samples. However, we noticed that Cluster 2 showed no differentially expressed marker genes. Thus, we performed comparative analysis of the QC metrics specific to this cluster (Figure S2B). This revealed significantly lower gene counts in cells assigned to Cluster 2, despite reduced mitochondrial content, suggesting that this cluster primarily contains low-quality cells, likely representing segmentation artifacts, and damaged cells, which could be excluded from downstream analysis.

Pathway analysis further resolved differences among neuronal subsets. Cluster 3 was enriched for genes associated with general neuronal identity and was composed predominantly of excitatory neurons, whereas Cluster 4 corresponded to inhibitory neuronal populations. The non-neuronal clusters corresponded to glial subpopulations and vasculature structures. Cluster 6 and 9 contained astrocyte populations, Cluster 10 comprised microglia, Cluster 11 represented oligodendrocyte precursor cells (OPCs), and Cluster 5 was enriched for vascular and endothelial cells. Clusters 7 and 8 were annotated as oligodendrocyte subtypes. Notably, Cluster 7 was enriched for FTH1+ oligodendrocytes, which have been described to provide a potential neuroprotective role in the context of neurodegeneration ^21^.

Finally, we evaluated the impact of preservation methods on the spatial transcriptomics data quality by comparing various QC metrics. These metrics included the median number of UMIs and genes per cell, median cell size (2 µm per cell) and mitochondrial content percentage (Supplementary Table 4). Overall, fresh-frozen tissues demonstrate higher per-cell quality and larger cell sizes, suggesting fewer segmentation artifacts. In contrast, FFPE samples showed lower mitochondrial content indicating reduced susceptibility of the remaining cells to sample processing through *10X Visium HD* protocol. While FFPE samples generally contained more cells than fresh-frozen samples, this difference may be attributable to variations in tissue cutting and final architecture.

### Benchmarking *Illumina* and *Ultima* sequencing platform performance for *Visium HD*

We compared the performance of Illumina NovaSeq X Plus and *Ultima Genomics UG 100* sequencing platforms for *Visium HD* libraries generated from FF human brain tissue across three selected samples. To minimize biological and tissue-processing effects, we prepared tissues using an optimized *Visium HD* protocol and split each final library into two aliquots for sequencing on each platform. After running the alignment pipeline, we benchmarked platform performance by comparing SpaceRanger-derived mapping quality combined with molecular and gene complexity metrics (Supplementary Table 6).

In the initial outputs, we sequenced the Ultima libraries substantially deeper than the corresponding Illumina libraries (on average, 685.3 million vs 289.0 million reads per sample, a 2.4-fold increase), resulting in higher sequencing saturation (88.43% in Ultima vs 80.10% in Illumina). The total genes detected were similar across platforms (16,868 vs 16,798), while probe-read base quality (Q30) was lower for Ultima (80.6% vs 97.5%). However, this reduction did not translate into lower downstream molecular quality. Read-mapping metrics were broadly similar between platforms, with Ultima showing a small increase in reads confidently mapped to the filtered probe set. Ultima libraries completely lacked reads half-mapped to the probe set, but these might be filtered out during the pseudo-pair regeneration process. Consistent with higher saturation, Ultima showed increased complexity per 8 µm bin, with the mean number of UMIs increasing from 80 to 108.3 (+35.4%) and the mean number of genes increasing from 73.3 to 87.5 (+19.4%). Mean mitochondrial content was essentially unchanged between platforms (5.08% for Illumina and 5.06% for Ultima), indicating that the increased Ultima library complexity did not originate from mitochondrial reads.

We downsampled Ultima libraries to match the Illumina read depth of each sample. After downsampling, read counts were effectively matched between platforms, and *Ultima* showed lower sequencing saturation than Illumina (77.8% vs 80.1% on average). Despite lower saturation, *Ultima* retained marginally higher molecular and gene complexity per 8 µm bin. The mean number of UMIs remained 10.3% higher for downsampled *Ultima* than Illumina (88.2 vs 80), and the mean number of genes remained 3.8% higher (76.1 vs 73.3).

Overall, these results demonstrate that the *Ultima UG 100* sequencing platform performs comparably to Illumina NovaSeq X Plus at matched sequencing depth for probe-based *Visium HD* libraries in fresh-frozen human brain tissue. Although *Ultima* showed lower Q30 values, the Space Ranger alignment metrics and per-bin QC suggest that this did not negatively affect overall data quality. These results show that this platform may offer a cost-effective strategy to increase feature and molecule recovery in probe-based *Visium HD* experiments, by enabling a deeper sequencing depth without increasing sequencing costs.

## Discussion

In this study, we evaluated and optimized the *10X Genomics Visium HD* workflow for spatial transcriptomics in human post-mortem brain tissue, focusing on both FF and FFPE preservation strategies. Our results show that, beyond the tissue preservation method, workflow execution plays a critical role in determining RNA preservation, probe capture efficiency, and overall library quality. We specifically address an important gap in the literature and provide a well-tested protocol has been specifically optimized to reduce the impact of RNA instability and workflow interruptions in FF and FFPE post-mortem brain tissue.

A major challenge for both tissue preservation methods is maintaining RNA integrity during cryosectioning and downstream handling ^22,23^. Our results in FF show that OCT embedding, required for proper sectioning, does not lead to major RNA degradation, but its impact is dependent on the initial RNA quality of the tissue ^22,24^. This is consistent with the variability observed between donors, where differences in baseline RNA integrity influenced downstream performance. These findings reinforce that FF tissue remains suitable for spatial transcriptomics but emphasize that both pre-analytical tissue quality and handling conditions are critical for success.

When applying the standard *Visium HD* workflow, including manufacturer’s recommended stop points, we observed suboptimal library quality, reflected by high Cq values and inconsistent fragment size distributions. This led to test our hypothesis that workflow interruptions, including long storage steps and repeated temperature changes, negatively affect RNA stability and probe capture efficiency. Although stop points are designed to provide flexibility, our data indicate that they are not well suited for FF and FFPE post-mortem brain tissue, where RNA is already degraded to some extent ^7,24^. This likely explains variability often observed across *Visium HD* studies using human brain samples, where technical limitations are not always explicitly addressed.

By optimizing the protocol into a continuous two-day workflow and extending probe hybridization from 16 to 20 hours, we observed a clear improvement in library quality. The lower Cq values and the consistent generation of fragments around the expected ∼250 bp size indicate enhanced probe capture and better preservation of RNA. These improvements were reproducible across consecutive tissue sections from the same donor, supporting the robustness of this approach. Overall, these findings highlight that reducing workflow interruptions and optimizing hybridization conditions are key factors for improving spatial transcriptomics performance in human post-mortem brain tissue. Moreover, they suggest that hybridization efficiency may represent a common limiting step across both FF and FFPE workflows, and that targeting this parameter can broadly improve data generation independent of preservation method.

An additional factor that likely contributed to the improved performance of the optimized workflow is the reduction of handling time during IF staining. The use of primary-conjugated antibodies would represent an effective strategy to minimize the number of incubation and washing steps, thereby reducing total processing time and limiting RNA exposure to suboptimal conditions. In our experience, strict adherence to the incubation times recommended in the *10X Genomics* protocol is also critical, as extending incubation periods, particularly during antibody staining, may increase the risk of RNA degradation. This is especially relevant for FF post-mortem brain tissue, where RNA is inherently unstable. Furthermore, maintaining tissue sections on ice whenever possible between experimental steps proved important to preserve RNA integrity throughout the workflow. Together, these considerations highlight that, in addition to global protocol design, careful control of staining processing conditions is essential to ensure optimal spatial transcriptomics outcomes.

In addition to protocol optimization, our sequencing saturation analysis provides insights into the relationship between sequencing depth and data quality. Although the recommended sequencing depth achieved high saturation levels, our results show that the number of detected genes and UMIs reached a plateau at significantly lower sequencing depths. Increasing sequencing depth beyond this point resulted in only modest gains in saturation and did not improve overall data quality. This indicates that library quality and probe capture efficiency, rather than sequencing depth, are the primary limiting factors in *Visium HD* experiments using FF post-mortem brain tissue. These findings have practical implications for experimental design and suggest that sequencing resources could be optimized once high-quality libraries are obtained.

While *Visium HD* technology offers whole-transcriptome spatial profiling and the ability to investigate cellular organization in complex tissues at unprecedented single-cell resolution ^25^, it is prone to analytical challenges driven by low transcript yields, reduced sensitivity, and barcode dissociation from individual cells. We were able to address those limitations by establishing an end-to-end QC and analysis pipeline that is agnostic to the tissue preservation method. We further demonstrated that this workflow is scalable across multiple donors, while maintaining the ability to extract coherent biological structures despite substantial sample heterogeneity.

A central methodological improvement in our pipeline was the shift from bin-based quantification to DAPI-based nuclear segmentation. Importantly, the resulting cells produced QC distributions comparable to those from 8-µm bins, while exhibiting a unimodal distribution of UMIs and detected genes at low counts, consistent with the exclusion of empty space. This supports nuclei segmentation as a practical strategy for improving the interpretability of *Visium HD* data without sacrificing “low-quality” transcripts relative to commonly used binning resolutions. While we demonstrate that fresh-frozen have higher quality compared to FFPE tissues from a data processing perspective (Supplementary Table 4), this finding may depend on tissue selection and requires additional samples for further validation.

Integration across samples remains a major analytical barrier for *Visium HD* data analysis. Even within a shared brain region, we observed minimal overlap in per-sample HVGs, indicating strong sample-specific biases that can propagate into noisy principal components and poorly resolved clustering (19). We elaborate that HVG selection can amplify technical effects rather than biological structure when transcript yields are low and heterogeneity is high. To address this, we implemented reference-based HVG selection using top marker genes derived from a large single-nucleus RNA-seq dataset (AUSBB cohort) and used these genes for scVI integration model training. Although reference HVGs overlapped only modestly with spatial HVGs, this strategy substantially improved cross-sample integration and produced coherent shared biologically relevant clusters. Additionally, spatial mapping of annotated cell types recapitulated expected tissue architecture, including clear gray/white matter demarcation, oligodendrocyte enrichment in white matter, and neuronal predominance in gray matter. Samples containing large vessels showed well-defined vascular structures populated by endothelial/vascular cells.

Importantly, the integration of IF imaging with spatial transcriptomics demonstrated that the protocol modifications did not alter the spatial distribution of AD pathology. The consistent detection of Aβ plaques and neurofibrillary tangles across conditions supports the compatibility of the optimized workflow with multimodal spatial analyses, which is essential for studying disease-associated molecular patterns in the human brain (20, 21).

New high-resolution spatial technologies, such as *Visium HD*, increase both technical and cost demands, prompting researchers to evaluate alternatives to standard Illumina sequencing ^28–30^. The *Ultima Genomics* platform has previously been benchmarked for single-cell RNA-seq analysis and demonstrated performance that is highly comparable to Illumina ^31^. Here, we compared the *Ultima Genomics UG 100* and Illumina NovaSeq X Plus platforms for probe-based *Visium HD* libraries from fresh-frozen human brain tissue. Despite lower Q30 scores, Ultima produced comparable alignment and QC metrics and retained marginally higher molecular complexity at matched sequencing depth. Given its lower sequencing cost, *Ultima* could be particularly beneficial when deeper sequencing is required to improve UMI and gene recovery in complex tissues such as the human brain.

This study has some limitations that should be considered. First, the optimization strategy was evaluated using a limited number of donors and focused primarily on specific cortical regions, which may not fully capture the biological and technical variability across different brain areas or disease stages. In addition, although improvements in library quality and sequencing efficiency were consistently observed, further investigation is required to determine how these gains translate to downstream biological analyses, including cell type resolution and detection of low-abundance transcripts. Another limitation is that our comparisons between FF and FFPE workflows were performed under optimized conditions, and additional studies are needed to assess performance across a broader range of tissue qualities and post-mortem intervals.

Additional limitations persist in the analysis of *Visium HD* transcriptomics data. Although our pipeline yields stable clusters that improve cell type annotation and capture spatial architecture, it does not fully address the missing spatial context in downstream analyses. As a result, spatial neighborhood information and tissue-domain organization may not be optimally captured. Segmentation-based approaches for brain tissues remain vulnerable to doublets and boundary errors ^32,33^, associated with densely packed regions. Nuclear expansion can also increase the probability of merging nearby cells, potentially inflating mixed profiles. Consistent with this limitation, we observed evidence that at least one cluster (Cluster 2) may be enriched with segmentation artifacts. Improved segmentation and cellular deconvolution tailored to spatial data informed by both imaging and transcriptomic signals will be necessary to further reduce these artifacts.

Future work should expand these analyses to larger cohorts, include additional brain regions, and explore integration with other spatial and multi-omics approaches to improve resolution and biological interpretation. Additionally, incorporating spatially aware models or explicit spatial clustering steps will be important to better leverage spatial resolution.

Overall, this study demonstrates that workflow optimization is essential for successful spatial transcriptomics in human post-mortem brain tissue. By removing stop points and extending hybridization time, we significantly improved probe capture efficiency and library quality in both FF and FFPE samples. Beyond data generation, our preservation-agnostic pipeline turns this heterogeneous material into a unified, interpretable resource. DAPI-based nuclear segmentation recovers single-cell resolution without sacrificing data quality, while sample-specific QC and reference-guided integration absorb technical variability to yield robust, reproducible cell-type annotation across samples and preservation methods. These results directly address the current gap in optimized workflows for human post-mortem brain, particularly FF preserved, and provide a practical approach that can be readily implemented to improve data quality and reproducibility in spatial transcriptomics studies of neurodegenerative diseases.

## Acknowledgements

We are grateful to the many donors, researchers, and staff who assisted with this study. We acknowledge the use of tissues procured by the National Disease Research Interchange (NDRI) with support from NIH grant U42OD11158. We thank the NRI-BBB, their staff, and the donors. Samples from the NRI-BBB were collected under The Ohio State University Institutional Review Board protocol#: 2020H0512. Tissues were received from the New South Wales Brain Tissue Resource Centre at the University of Sydney, which is supported by the University of Sydney and National Institute of Alcohol Abuse and Alcoholism of the National Institutes of Health under Award Number R28AA012725, the content is solely the responsibility of the authors and does not represent the official views of the National Institutes of Health.

Research reported in this publication was supported by The Ohio State University Comprehensive Cancer Center and the National Institutes of Health under grant number P30 CA016058. We thank the Genomics Shared Resource at The Ohio State University Comprehensive Cancer Center, Columbus, OH for quality control assistance. We would also like to thank Kara Corps, Nicholas Sweeney, and Pearlly Yan for assistance with imaging the FFPE slides, and Ishrat Jahan for general laboratory assistance.

Data generation was supported by the following NIH grants: U01AG07246403, R01AG07401201, RF1NA142335, R61NA138655, R01AG075092 and R56AG06776401. Additional funding to The Ohio State University and the researchers on this project was provided by the Koch Alzheimer’s Research Fund. We also thank the donors and their families for their generous brain donations to help further our knowledge of Alzheimer’s disease.

## Code and data availability

Custom code used to analyze the spatial transcriptomics data is available at https://github.com/HarariLab/2026_visium_hd_methodology. Single nuclei RNA sequencing reference dataset located on AD Knowledge Portal at Synapse ID syn69831812.

## Authors contribution

A.C.B., E.A., H.T.N.H., G.T.S., and O.H. contributed to the study conception and experimental design, data analysis, data interpretation, intellectual input, and manuscript preparation and revision. A.C.B. processed FF samples and optimized the spatial transcriptomics protocol. H.T.N.H. processed FFPE tissue samples and optimized the spatial transcriptomics protocol. E.A. performed the bioinformatics analyses and data integration. K.A. assisted with bioinformatics and data analysis. A.C.B., E.A. and K.A. prepared all figures, tables, and supplementary materials. A.C.B. and S.A. performed immunofluorescence staining and imaging of FF brain tissues. N.S. and T.Y.K. performed immunofluorescence staining and imaging of FFPE brain tissues. K.L.B. collected the FF brain tissues. I.D.D.S., R.D.A., H.F. and G.M. contributed with the study design. O.H and G.T.S secure funding for the project. All authors reviewed, revised, and approved the final version of the manuscript.

---

**Supplementary Figure 1**: Effects of sequencing depth on QC metrics. (A) Association between the mean number of UMIs and genes and changes in sequencing depth. Panels are separated by aggregates’ resolutions – 2, 8 and 16 µm bins, segmented cells. (B) Additional plots (see Figure 7B) showing the association between sequencing depth and the total number of expressed genes for 2 and 16 µm bins. For (A) and (B), the target levels represent the 10X minimal recommendations for sequencing depth.

**Supplementary Figure 2**: QC analysis. (A) Violin plots comparing distributions of the number of UMIs and mitochondrial content percentage between different aggregates – 8 µm bins and segmented cells. (B) Heatmap of hierarchical clustering of HVGs between samples and the snRNAseq reference dataset, based on the Jaccard similarity index. No groups with significant overlap were detected. (C) Cell density plots on an integrated UMAP, with individual sample panels. Only sample ffpeAD3_S1 demonstrates remaining bias.

**Supplementary Figure 3**: Cell type associations for detected unsupervised clusters. (A) Dot plot of expression of known cell-type markers per cluster. (B) Dot plot of cluster markers identified by differential expression analysis. (C) Gene set enrichment analysis (GSEA) with the top 3 terms per cluster. Gene sets for GSEA were obtained from the “Azimuth_Cell_Types_2021” database.

**Supplementary Figure 4**: QC metrics for each detected unsupervised cluster. (A) Additional spatial plots (Figure 7F) with annotated cells. (B) Violin plots showing the distributions of gene counts and mitochondrial content percentage per cluster. Cluster 2 has a significantly lower number of genes than other clusters and a low mitochondrial content.

