## Supplementary figures and images for "An end-to-end framework for single-cell-resolution, whole-transcriptomic spatial profiling in post-mortem human brain"

### Figure S1

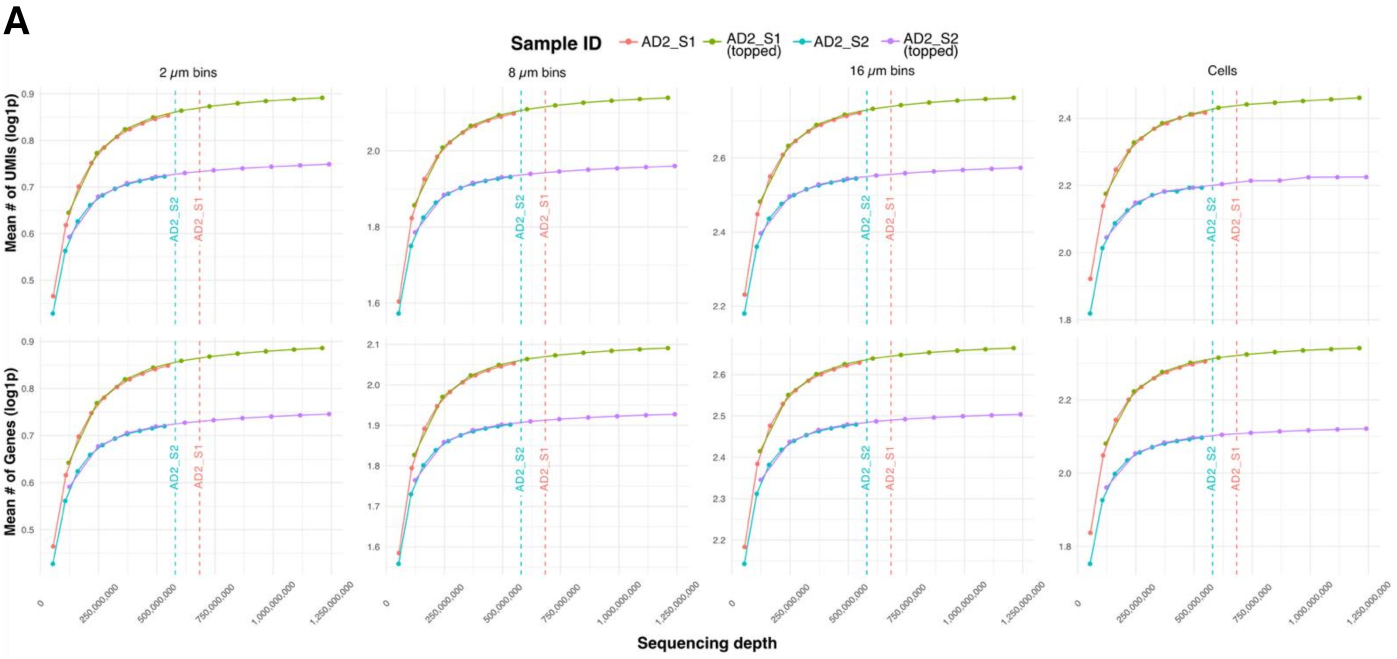

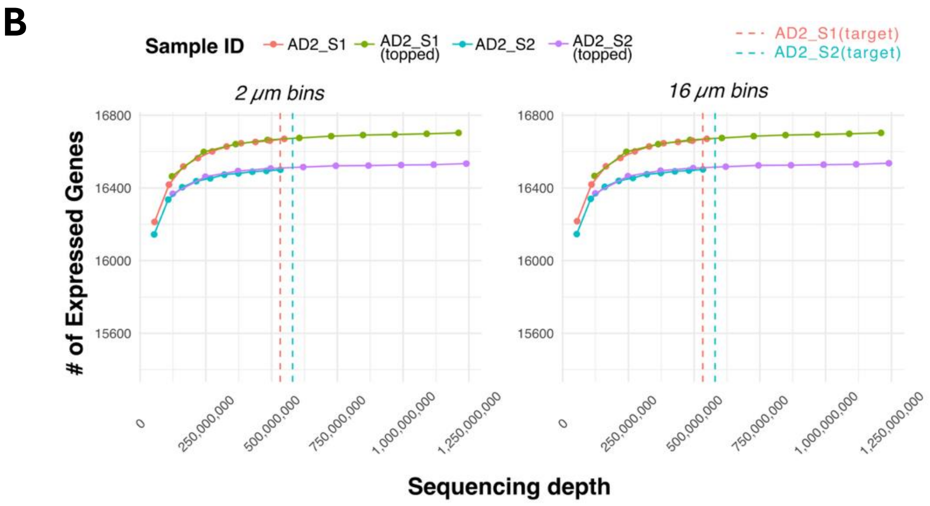

### Figure S2

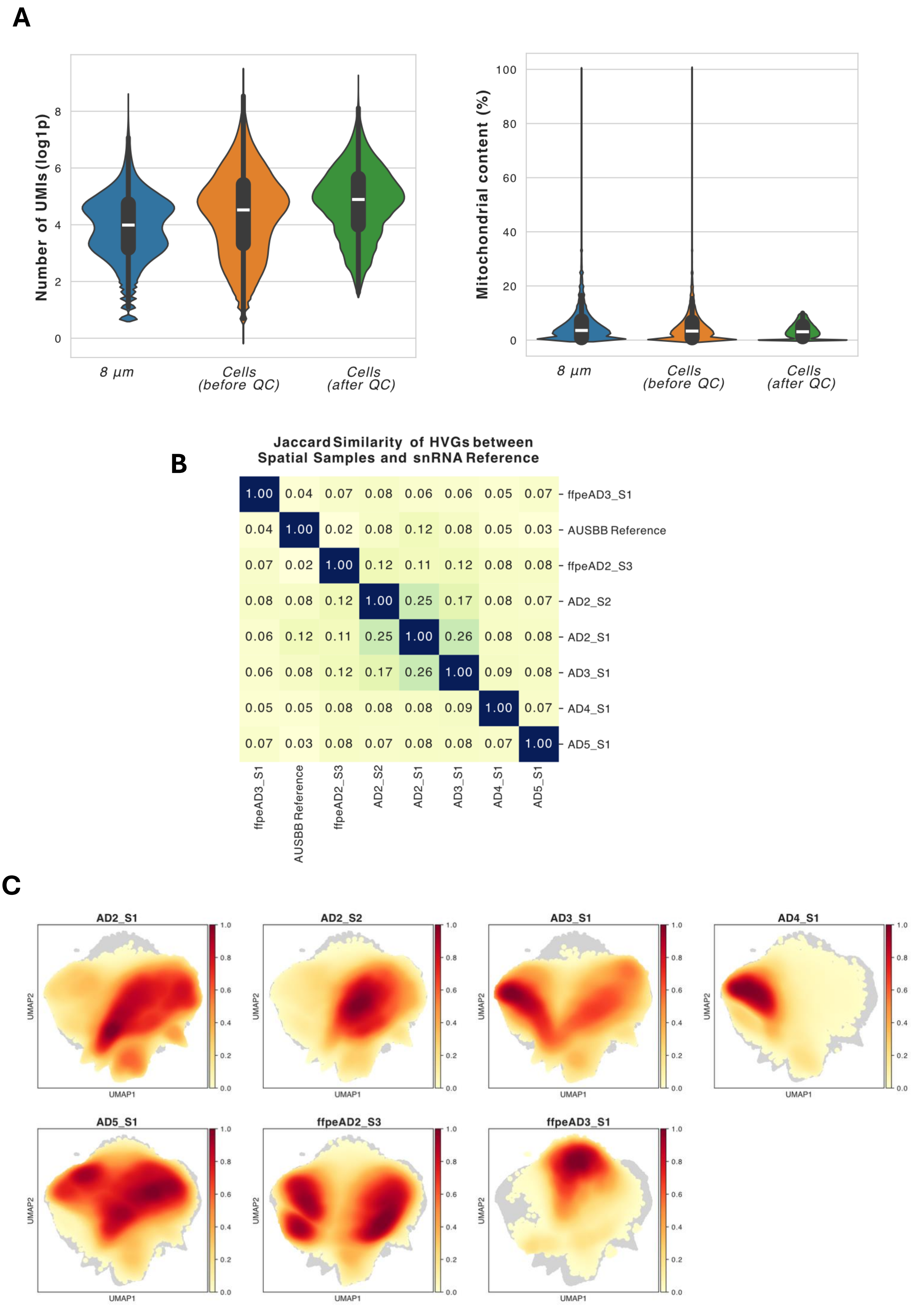

### Figure S3

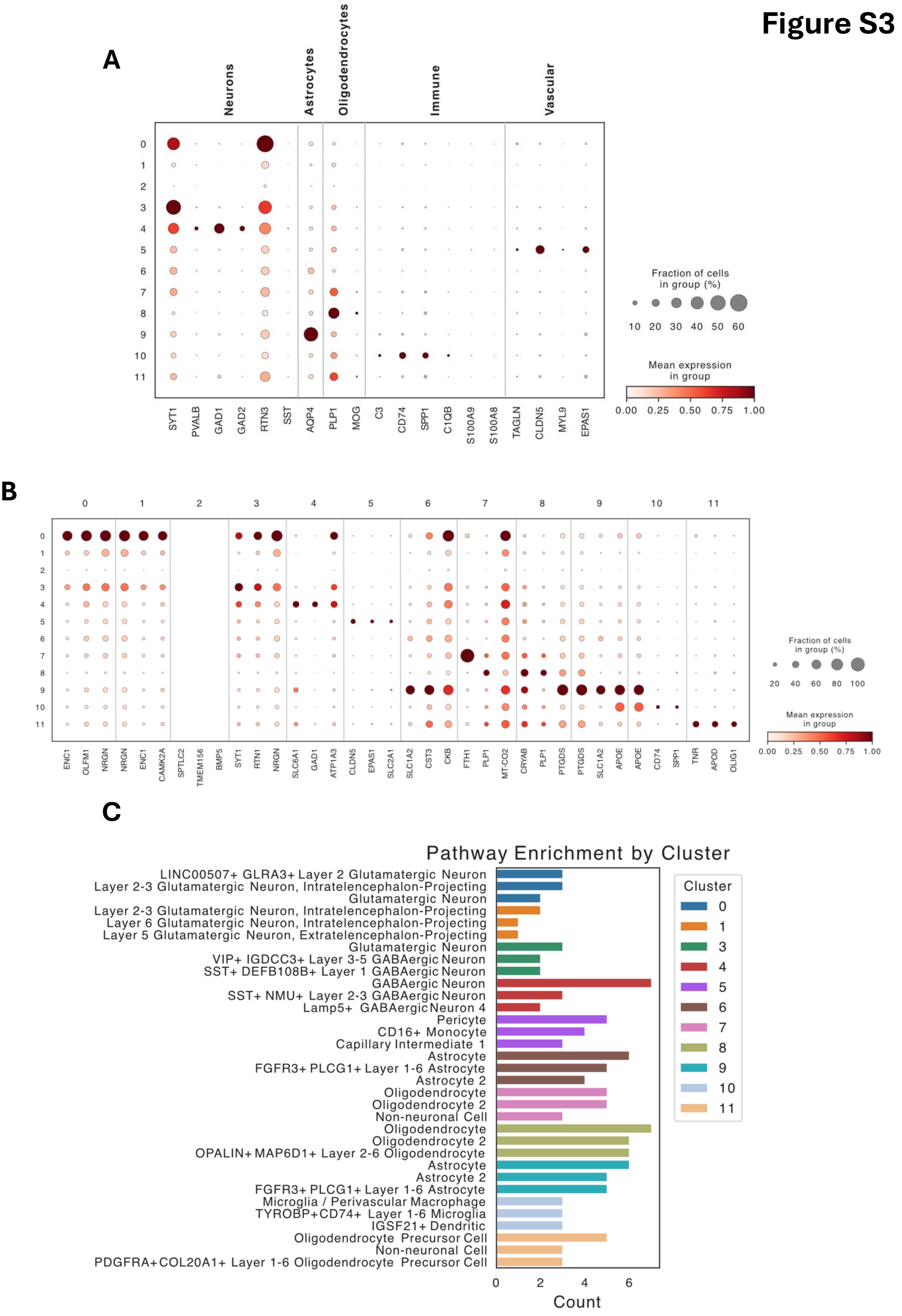

### Figure S4

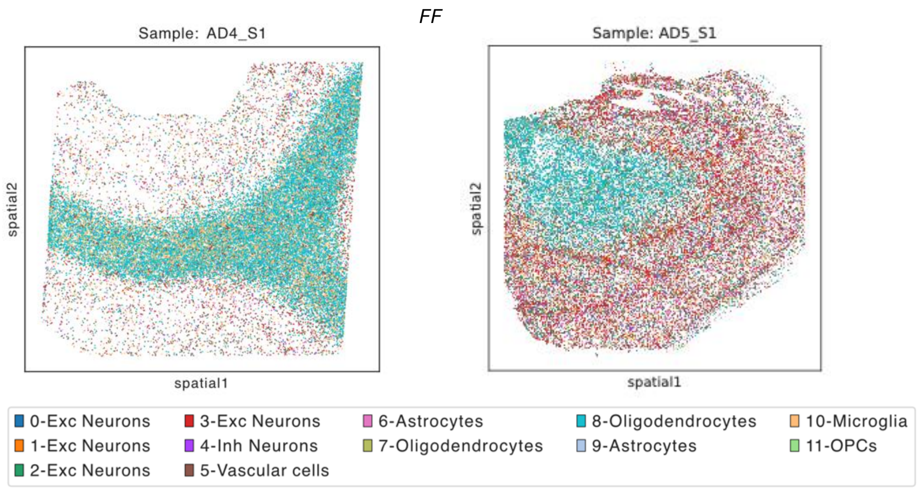

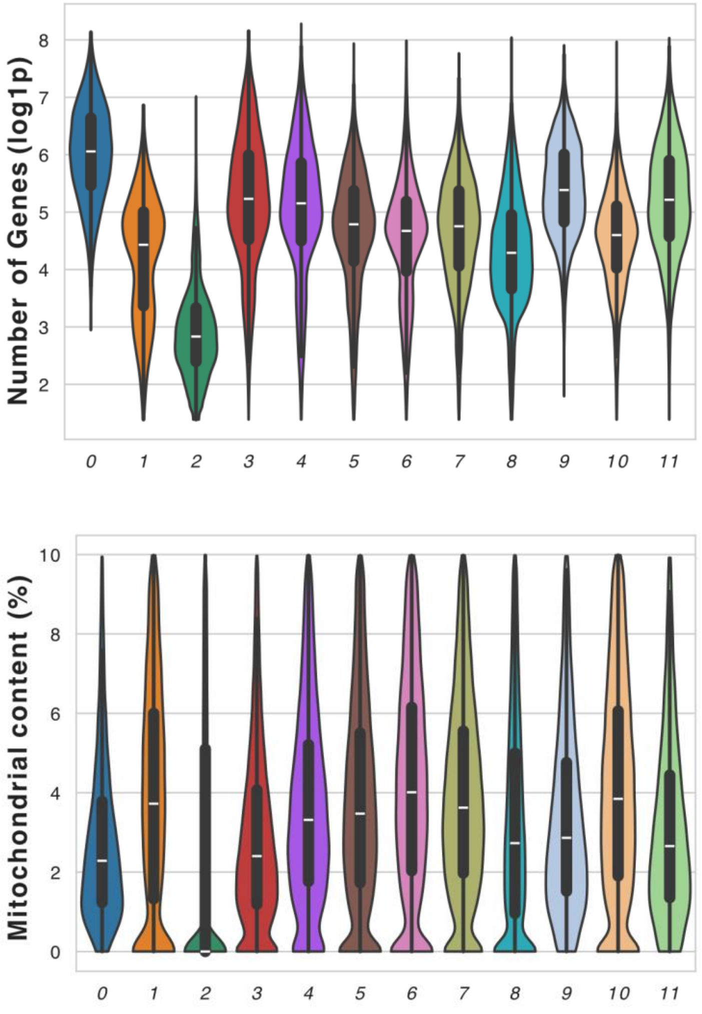
